# Multiplexed Profiling of Single-cell Extracellular Vesicles Secretion

**DOI:** 10.1101/459438

**Authors:** Yahui Ji, Dongyuan Qi, Linmei Li, Haoran Su, Xiaojie Li, Yong Luo, Bo Sun, Fuyin Zhang, Bingcheng Lin, Tingjiao Liu, Yao Lu

## Abstract

Extracellular vesicles (EVs) are important intercellular mediators regulating health and disease. Conventional EVs surface marker profiling, which was based on population measurements, masked the cell-to-cell heterogeneity in the quantity and phenotypes of EVs secretion. Herein, by using spatially patterned antibodies barcode, we realized multiplexed profiling of single-cell EVs secretion from more than 1000 single cells simultaneously. Applying this platform to profile human oral squamous cell carcinoma (OSCC) cell lines led to deep understanding of previously undifferentiated single cell heterogeneity underlying EVs secretion. Notably, we observed the decrement of certain EV phenotypes (e.g. ^CD63+^EVs) were associated with the invasive feature of both OSCC cell lines and primary OSCC cells. We also realized multiplexed detection of EVs secretion and cytokines secretion simultaneously from the same single cells to investigate multidimensional spectrum of intercellular communications, from which we resolved three functional subgroups with distinct secretion profiles by visualized clustering. In particular, we found EVs secretion and cytokines secretion were generally dominated by different cell subgroups. The technology introduced here enables comprehensive evaluation of EVs secretion heterogeneity at single cell level, which may become an indispensable tool to complement current single cell analysis and EV research.

**Significance:** Extracellular vesicles (EVs) are cell derived nano-sized particles medicating cell-cell communication and transferring biology information molecules like nucleic acids to regulate human health and disease. Conventional methods for EV surface markers profiling can’t tell the differences in the quantity and phenotypes of EVs secretion between cells. To address this need, we developed a platform for profiling an array of surface markers on EVs from large numbers of single cells, enabling more comprehensive monitoring of cellular communications. Single cell EVs secretion assay led to previously unobserved cell heterogeneity underlying EVs secretion, which might open up new avenues for studying cell communication and cell microenvironment in both basic and clinical research.

## Introduction

Extracellular vesicles (EVs), including exosomes, microvesicles, etc., are critical components in cellular microenvironment regulating intercellular communications and transferring biology information molecules like cytosolic proteins, lipids, and nucleic acids(1-6). Due to its relatively stable duration in circulation system, it has been used as noninvasive diagnostic markers for disease progression(7-9) or therapeutic treatments (2, 3). Detection and stratification of extracellular vesicles is crucial to increase our understanding of EVs and bring new applications in biomedicine, such as theirsizes(10), morphologies(11) and molecular compositions(12, 13). Among different molecular components involved in EVs functionalities, proteomic surface markers provide direct targets for intercellular communication mediated by EVs(14, 15). Thus, varieties of methods have been reported for profiling protein markers on EVs’ surface, such as ELISA(16), western blotting(15), flow cytometry(10), imaging(17), etc., from population of EVs(15) down to single vesicle level(10, 17). However, these measures are still at population cells level, which averaged EVs secretion from different cellular sources and obscured cell to cell heterogeneity in quantity/phenotypes of EVs secretion and their related functions (18-22). Nanowell-based (23, 24) and tetraspanin-based pH-sensitive optical reporters(25) for single cell EVs secretion assay have been developed to address the need, however with limited proteomic parameters (≤2) for EVs from each single cell, which is not sufficient to dissect EVs secretion heterogeneity comprehensively. A technology that can profile an array of surface markers on EVs from large numbers of single cells is still lacking and will help addressing a host of important biological questions ranging from inter and intra-tumor diversity, to cell-cell communication network and be of great value to clinical applications like personalized diagnostics and medicine.

Herein, we demonstrated a microchip platform for multiplexed profiling of single cell extracellular vesicles secretion to address the critical need for new technologies to dissect the communication spectrum of tumor cells mediated by EVs. The multiplexed profiling was realized with antibodies barcode, which is a reliable, reproducible technology previously adopted for blood testing(26), single cell proteomic analysis(27-31), and immunotherapy monitoring(32-34). We applied the platform to profile human oral squamous cell carcinomas cell lines and patient samples, which revealed previously unobserved secretion heterogeneity and identified the decrement of certain EV phenotypes (e.g. ^CD63+^EVs) were associated with the invasive potential of both OSCC cell lines and primary OSCC cells. Besides, we also realized simultaneous profiling of EVs secretion and cytokines secretion from the same single cells for deep understanding of cellular organizations and uncovering the correlation between different types of intercellular communication mediators.

## Results

### Platform for multiplexed profiling of single-cell EVs secretion

The platform to realize multiplexed single-cell EVs secretion detection (**Figure 1A**) is modified from previously reported devices (31), which combines two functional components: a high density microchambers array and spatially resolved antibodies barcode glass slide. High density microchambers array (**Figure S1A**) accommodates 6440 identical units for isolating and concentrating EVs secreted from exactly a single cell (each microchamber: width 40μm, length 1440μm, depth 30μm). The volume of each microchamber is around 1.7nL, which corresponds to 5×10^5^ cells/mL cell density, comparable to the cell density typically used in bulk experiments. Due to the drastic decrease in liquid volume, the concentration of detection targets will be concentrated as much as 10^3^−10^5^ times compared with population measurements (1.7nL vs 10-200μL), which ensured high sensitivity detection. We designed and fabricated accompanying microchip with highly parallel microchannel array to pattern spatially resolved antibodies barcode onto poly-L-lysine glass slide, which can accommodate up to 9 different antibodies (with each antibody stripe 40μm in width) for multiplexed profiling (**Figure S1B**, the consumption of each antibody for patterning is only 3μL volume at 250μg/mL). The antibodies patterning can be finished within 4hrs with excellent uniformity (fluorescein isothiocyanate labelled bovine serum albumin coating: C.V. <5% in 2cm x 5.5cm area, **Figure S2**).

**Figure 1.**
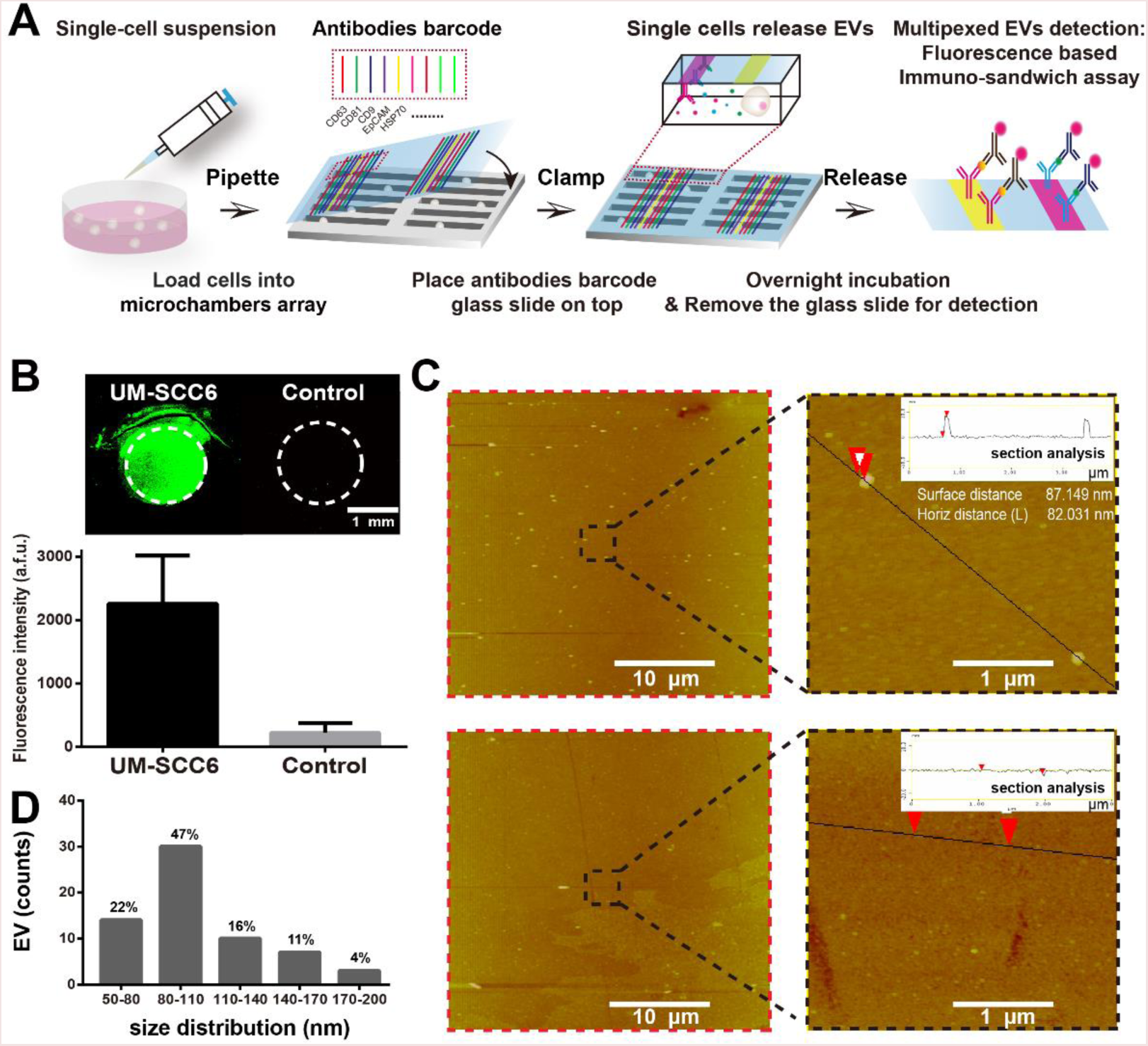
Platform for multiplexed profiling of single-cell extracellular vesicles secretion. A, Schematic illustration of the workflow for multiplexed profiling of single-cell extracellular vesicles secretion. Images of two functional components were shown in **Figure S1**. B, EV detection results on anti-human CD81 antibody coated spot with UM-SCC6 cells conditioned medium and control sample: blank cell culture medium supplemented with 10% ultra-centrifuged FBS. C, AFM characterization of fluorescence detection regions showing in B. D, Size distribution of extracellular vesicles captured on the anti-CD81 antibody coated surface.

We verified antibody based EVs capture/detection principle at bulk level (**Figure S3**). Antibodies targeting human CD81 and CD63 for EVs were used to form detectable immuno-sandwich, both of which are tetraspanins highly expressed in EVs for reliable EVs marker proteins (14, 15). This double positive detection strategy based on different epitopes recognition can eliminate the crosstalk from soluble molecules to ensure the detection specificity (16). We obtained positive fluorescence signals with conditioned medium from human oral squamous carcinoma cells (UM-SCC6) (**Figure 1B**). Atomic Force Microscope (AFM) characterization confirmed the fluorescent signals were from EVs (**Figure 1C**). The diameter of the captured particles ranged from 50nm to 200nm, suggesting EVs captured covered both exosomes (size: 50–150nm) and microvesicles(size: 100–1000nm) (**Figure 1D**). Consistent with fluorescence results, we didn’t capture any particles from exosome depleted cell culture medium sample. We further confirmed multiple EVs can be profiled on micrometer sized antibody stripes (**Figure S4**), demonstrating the feasibility to use antibodies barcode for multiplexed EVs detection.

### Multiplexed single-cell profiling reveals complex heterogeneity underlying EVs secretion

We then used the platform to profile the EVs secretion with human oral squamous cell carcinoma (SCC25) to assess its single cell detection sensitivity (35). 40,000 cells (200μL at 2×10^5^cells/mL density) was pipetted directly onto hydrophilic microchamber array (oxygen plasma treated). When enclosed by putting antibodies barcode glass slide on the top, more than 1000 single cells (1386±276, n=6) were constantly obtained, ensuring high-throughput analysis and statistical significance. The proteomic parameters for EVs surface marker profiling used in this study includes CD63, CD9, CD81, EpCAM and HSP70. With the combination of surface markers used here, the EVs captured from the same single cells can be further categorized into five subgroups: ^CD63+^EV, ^CD9+CD63+^EV, ^CD81^+^CD63+^EV, ^Epcam+CD63+^EV
and ^HSP70+CD63+^EV. A representative fluorescence detection result from SCC25 cells was shown in **Figure 2A**, from which we observed fluorescent positive square spots intersecting CD63/CD81/CD9 antibodies barcode with signal-to-noise ratio (SNR)≥3, demonstrating EVs with different surface proteins from the same single cells were reliably detected.

**Figure 2.**
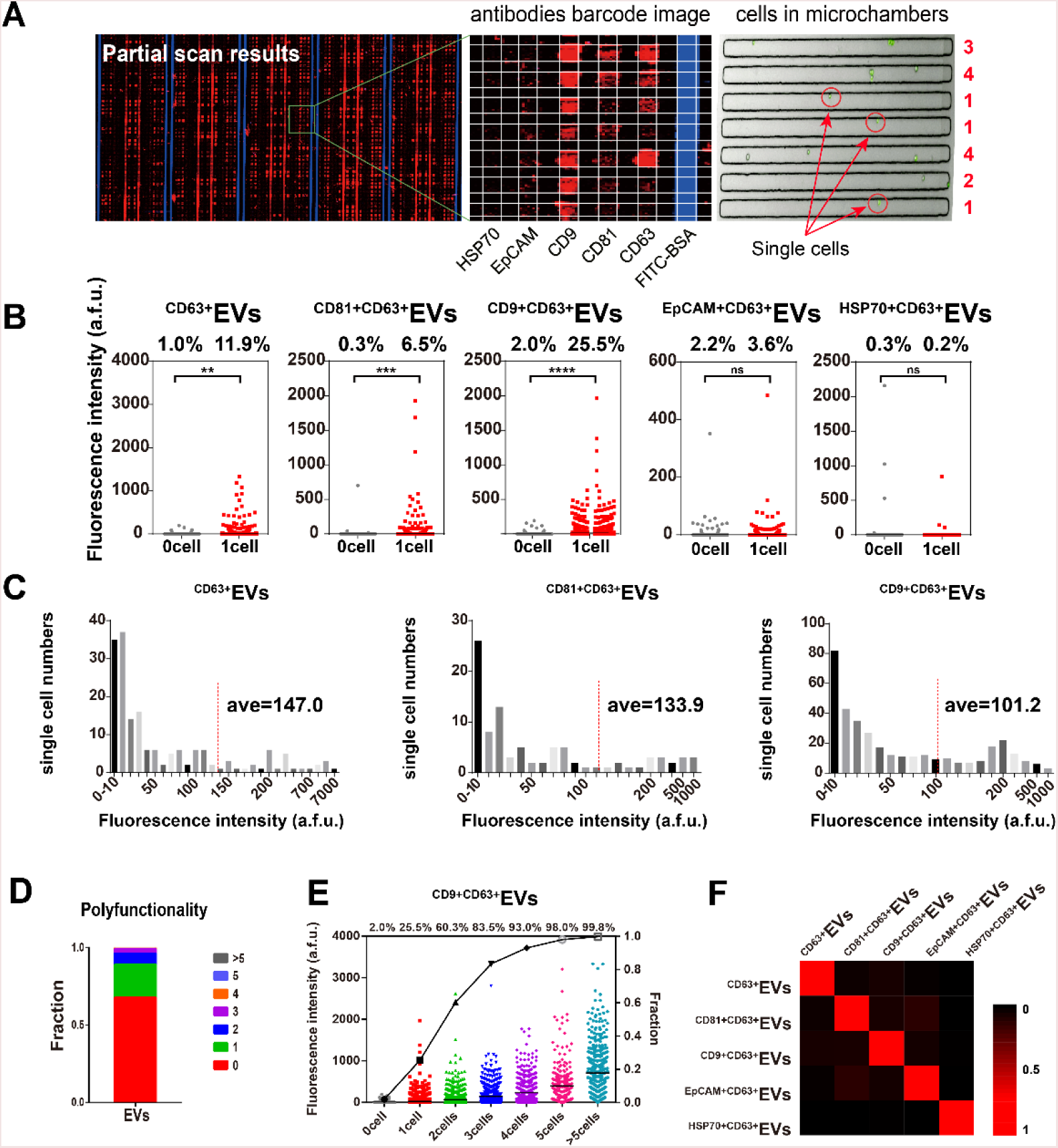
Multiplexed profiling of SCC25 single-cell EVs secretion revealed previously unobserved secretion heterogeneity. A, Representative images showing the raw data of multiplexed single-cell EVs profiling, including fluorescence detection images (partial and enlarged), corresponding cells in microchambers (red circled are single cells). B, Scatter plots showing 5-plexed EVs secretion profiling from SCC25single cells (n=1264, 1417 for zero cells and single cells respectively). The fluorescence intensity and secretion frequency were normalized with the average plus 2SD of the zero cell data. C, Histograms showing secretion intensity distribution of different EVs (^CD63+^EVs, ^CD81+CD63+^EVs and ^CD9+CD63+^EVs). D, Poly-functionality analysis of SCC25 single cells (fraction of cells to secrete multiply EVs simultaneously). E, Scatter plots showing the change of ^CD9+CD63+^EVsecretion frequency with the increased number of cells per microchamber. F, Heatmap showing EV-EV correlation in single cells.

After single cell data were normalized based on mean fluorescence intensity plus 2 times standard deviation (SD) of all zero-cell microchambers as thresholds to define positive secretion events (31), SCC25 single cell EVs secretion results were presented as scatter plots in **Figure 2B**, which provide direct insights to understand EVs secretion heterogeneity, i) not all cells can secrete EVs, for example, only around 6.2% cells secreted ^CD81+CD63+^EVs; ii) intensity distribution within these EVs secreting cells revealed that a very small number of cells can secrete ~10 times more than averaged secretion, indicating the presence of outliers or “super EV secretors” within cell population (**Figure 2C**); iii) cells secreted EVs with preference within different surface markers, for example, around 23.5% SCC25 cells secreted ^CD9+CD63+^EVs,IISr7Q I CD63 I while we could barely see +EVs secretion at single cell level; iv) a small fraction of SCC-25 cells could secrete multiple cytokines or EVs simultaneously (**Figure 2D**), for example, only ~2.7% single cells can secret EVs with more than three different combinations of surface markers at the same time, further confirming the presence of “super EV secretors” within cell population. Collectively, these observations presents the complex heterogeneity underlying EVs secretion, which is difficult to profile with population measurements. Interestingly, we found the percentage of cells with positive EVs secretion would increase with more cells in each microchamber (**Figure 2E**), suggesting EVs secretion is also mediated with paracrine signaling, which is in agreement with other report(36). We also saw that these EV phenotypes were weakly correlated via linear regression analysis of the correlation coefficient between EVs (**Figure 2F**).

### Decreased single cell EVs expression in invasive tumor cells

We then applied the platform to profile EVs derived from tumor cells with different migratory properties at single cell level to uncover the correlation between EVs secretion and cell’s invasive behavior. A subgroup of UM-SCC6 cells with high invasion behavior in matrigel matrix (named as UM-SCC6M) were obtained by three rounds of isolation of invasion front of UM-SCC6 cells in an H shaped microfluidic chip (**Figure 3A**). Detailed isolation procedures has been reported previously(37). To dissect the multidimensional spectrum of intercellular communications, here we profiled 5-plexed EVs secretion (CD63, CD9, CD81 and EpCAM, HSP70) along with 3-plexed proteins secretion (IL-6, IL-8, MCP-1) simultaneously from each single cells to provide direct correlation between different types of intercellular messengers (**Table S1**). Titration tests with recombinant proteins and antibodies crosstalk tests were completed to validate technical validity (**Figure S5&S6**). Interestingly, we found UM-SCC6M cells, which is active in invasion, were less active in secretion for both EVs and proteins, compared with UM-SCC6 cells (**Figure 3B**). Specifically, 12.3% of UM-SCC6 single cells secreted ^CD63+^EVs, while 4.3% of UM-SCC6M single cells secreted ^CD63+^EVs; 10.9% UM-SCC6 single cells secreted ^CD9+CD63+^Ev vs 0.9% for UM-SCC6M single cells; 10.6% of UM-SCC6 single cells were positive in IL-8 secretion, while only 5.7% of UM-SCC6M single cells secreted IL-8. Previous studies have demonstrated tetraspanins CD63, CD9 are metastasis suppressors, highly expressed in the early stages of different cancers (e.g. melanoma(38, 39), carcinoma(40, 41)) and decreased in advanced stages. Our results showed the CD63, CD9 expression were also decreased on tumor cell derived EVs surfaces when cells are in invasive state, which has never been observed previously at single cell level. We also saw the similar trend in cell population assay (**Figure 3C**) and a reasonable level of correlation between single-cell results and cell population measurements (**Figure 3D,** Pearson r = 0.76, P < 0.05), despite significant differences in assay conditions between them.

**Figure 3.**
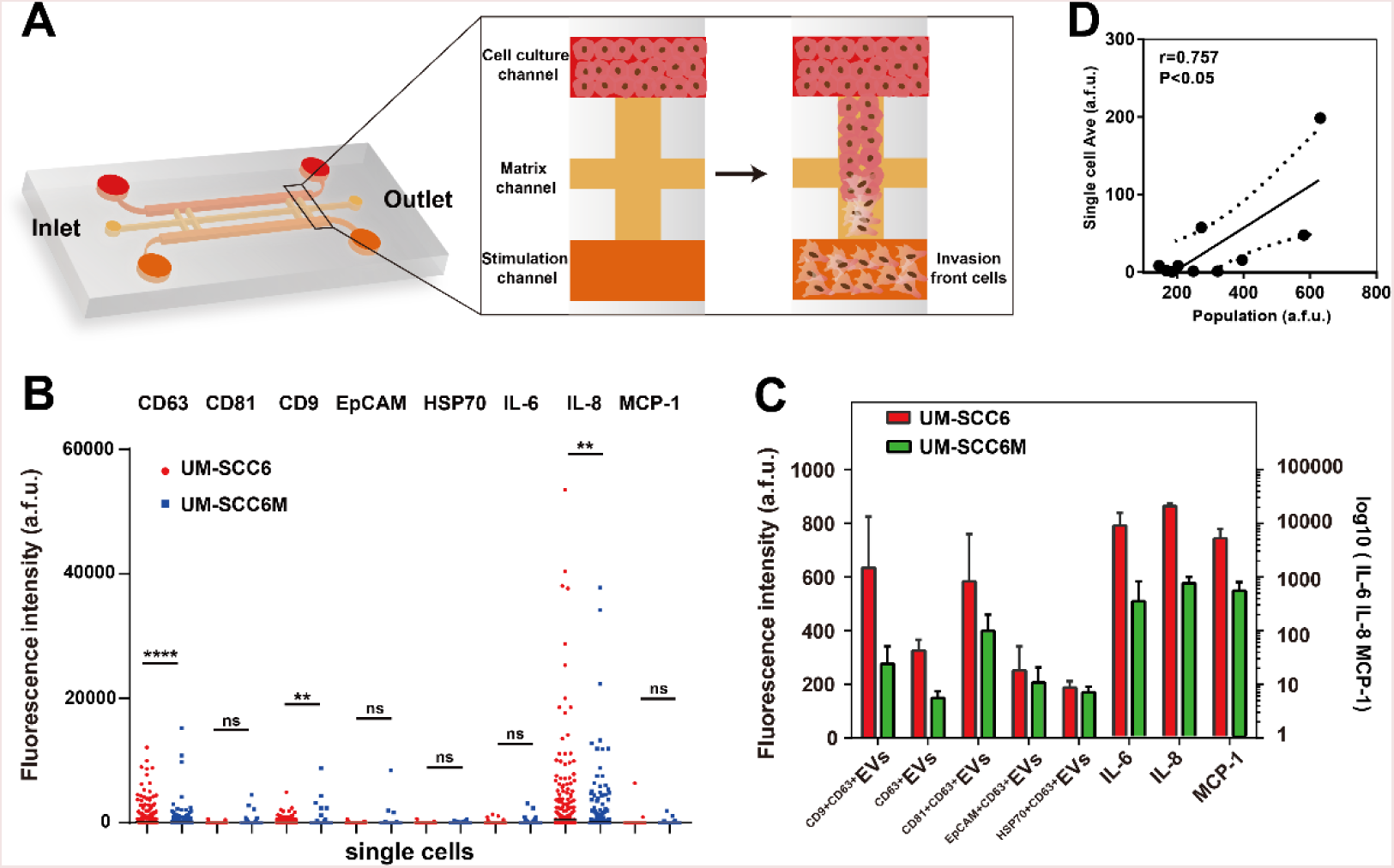
Single cell EVs secretion in invasive tumor cells. A, Illustration of the isolation of invasion front of UM-SCC6 cells in a microchip. B, Comparative analysis of secretion frequencies between UM-SCC6 (n=1263) and UM-SCC6M (n=1512) single cells. C, 8-plexed secretion profiles from cell populations. D, Correlation of EV secretion levels between single-cell averages and cell population measurements (the dashed line showed the 95% confidence interval).

### Multiplexed profiling of single-cell secretion of OSCC patient samples

To further demonstrate potential applications of our single cell analysis platform towards clinical samples, we profiled three primary ex vivo tissues from oral squamous cell carcinoma (OSCC) patients to discern metastatic tumor derived EVs associated with EVs secretion (**Table S2**). The fresh OSCC tumor tissues from surgery were disassociated, purified into primary tumor cell suspensions and verified with epithelial malignancy marker pan Cytokeratin immunostaining (42) (**Figure 4A**). Though three tumor samples were all identified as high differentiation grade (**Table S2 & Figure S7**), patient 1&2 were diagnosed as metastatic, while patient 3 was non-metastatic. All three patients exhibited similar secretion signatures as OSCC cell lines, for example, they were relatively strong in ^CD9+CD63+^EVs and IL-8 secretions, while attenuated in ^EpCAM+CD63+^EV and ^HSP70+CD63+^EV secretions. Notably, we observed ^CD63+^EVs and ^CD81+CD63+^EVs secretion in metastatic patients 1&2 was significantly decreased, compared with non-metastatic patient 3 (**Figure 4B**), which confirmed our observation from UM-SCC6 cell line. While ^CD9+CD63+^EVs didn’t show significant differences between patients, suggesting the heterogeneity between cell lines and primary cells. Unsupervised clustering further resolved tight correlation between patient 1 and 2 in EVs secretion pattern (Spearman correlation r_1_,_2_=0.975, r_1_,_3_=0.667, r_2_,_3_=0.6) (**Figure 4C**).

**Figure 4.**
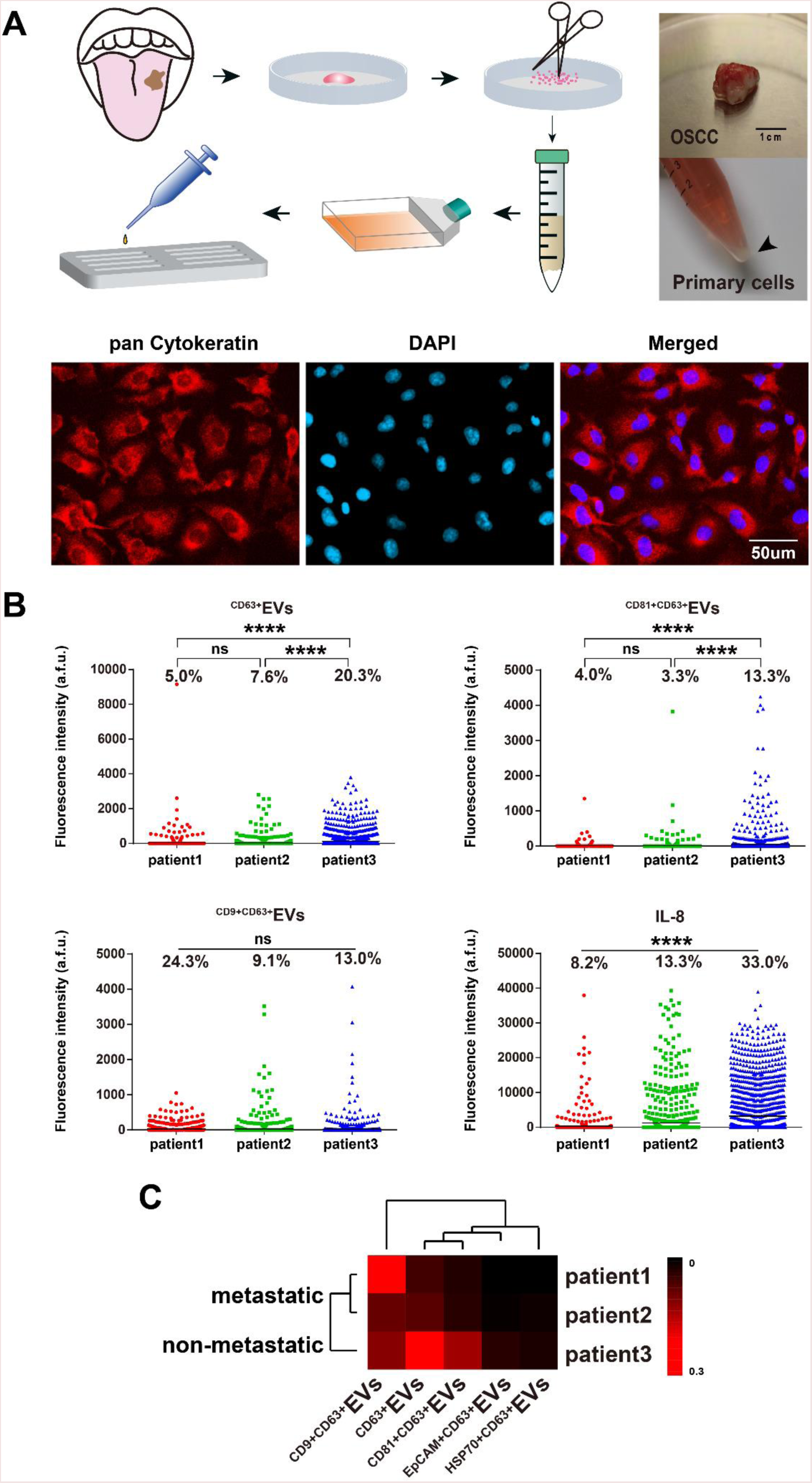
Single-cell secretion analysis of primary tumor cells from OSCC patients. A, Overview of disassociating patient surgery specimens into primary tumor cells suspension (verified with pan CK immunostaining) and its follow-on procedures to apply primary cells to microchamber array for single cell analysis. B, Comparative analysis of individual secretions (^CD63+^EV, ^cD81+CD63+^EV, ^eD9+eD63^+EV and IL-8) among patients. (ns>0.05, ***< 0.0005 by t test, n=974, 1351 and 1801 respectively for three patients). C, Clustering of three patient samples based on secretion frequencies of all EVs parameters.

### Single-cell secretions functional phenotyping

We then mapped all the single cell data from OSCC patient samples using viSNE(43), which is based on the t-Distributed Stochastic Neighbor Embedding algorithm, to reveal their functional organizations (**Figure 5A**). We saw primary cells from each patient gave rise to three structured clusters: Group 1 is mainly distinguished with EVs secretion, like ^CD9+CD63+^EVs and ^CD63+^EVs; Group 3 dominated proteins secretion, mainly for IL-8; while Group 2 accommodates both EVs secretion and proteins secretion, but with much attenuated frequency. Interestingly, we observed similar functional organizations in OSCC cell lines (**Figure S8&S9**), demonstrating the functional architecture of population cells is relatively stable across both cell lines and primary cells (28).

**Figure 5.**
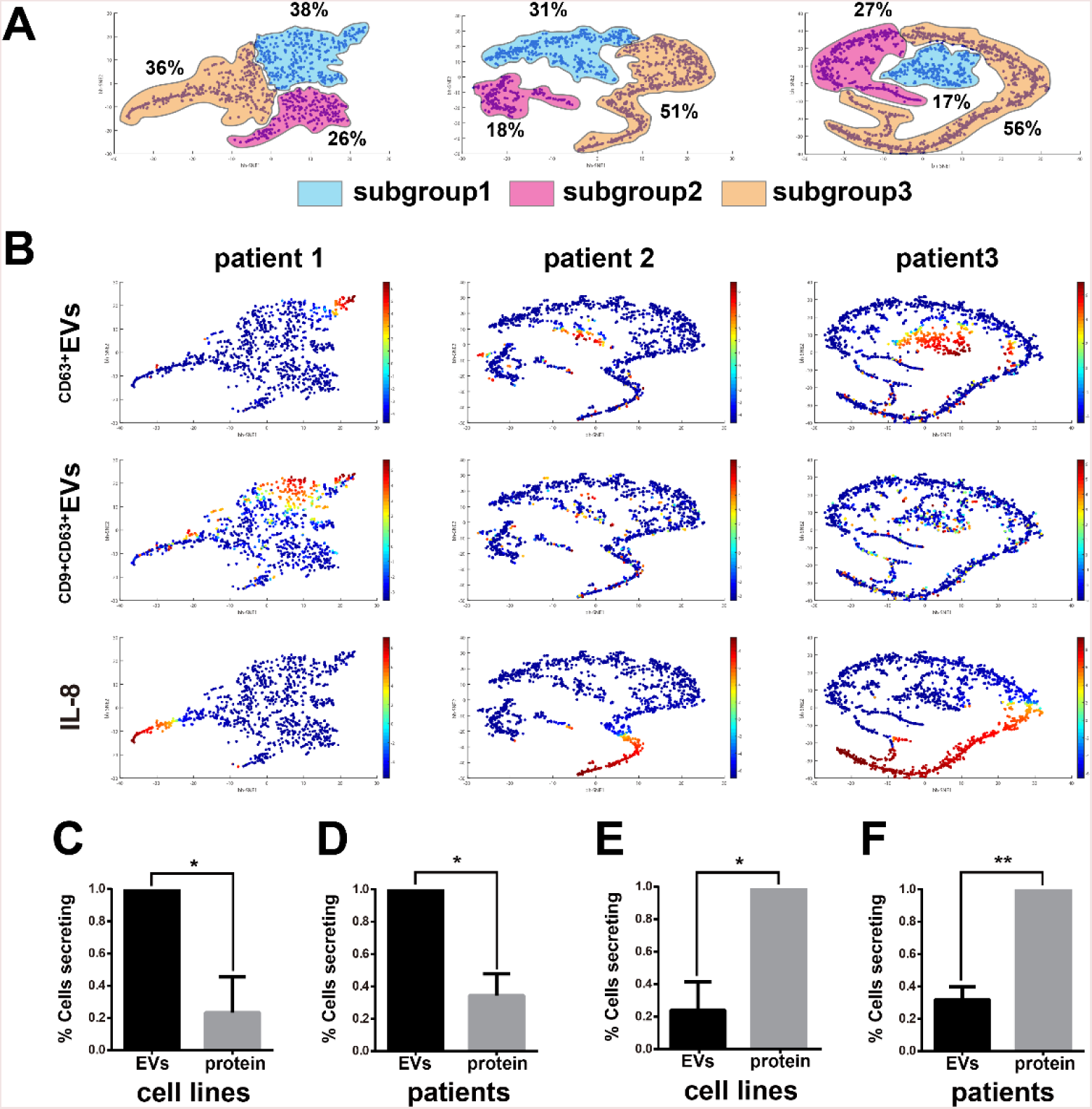
Multidimensional single cell secretion analysis delineated OSCC cellular functional organization. A, Visualized clustering analysis with viSNE revealed three functional subgroups in all patient samples. B, Distribution of individual secretors (^CD63+^EV, ^CD9+CD63+^EV and IL-8) in patients derived viSNE plots. C-D, Probability of EVs secreting cells to secrete proteins simultaneously. (C: cell lines, D: patients); E-F, Probability of proteins secreting cells to secrete EVs simultaneously. (E: cell lines, F: patients) simultaneously. (*< 0.05, **< 0.005 by paired t test, n=3).

From the viSNE maps (**Figure 5B & Figure S8-12**), we further observed the cells positive with protein secretions were less likely to secrete EVs simultaneously, suggesting the EVs and proteins secretion were generally dominated by different cell subsets within population. To confirm this finding, we calculated the conditional probability of EV positive cells to secrete proteins and found EV+ cells were significantly less likely to secrete proteins in both cell lines (SCC25, UM-SCC6&UM-SCC6M) (p=0.027 by paired t test) and patients (p=0.014) (**Figure 5C&D**). Likewise, protein+ cells were significantly less likely to secrete EVs simultaneously in both cell lines (p=0.017) and patients (p=0.005) (**Figure 5E&F**).

## Discussion

Tumor derived EVs play important roles in tumor metastatic processes(2, 6, 7), which make it vital to obtain more detailed information from these vesicles. However, these EVs were never characterized comprehensively at single cell level, due to the lack of available tools. This study introduced, for the first time (to the best of our knowledge), an antibody barcode based platform for high throughput, multiplexed profiling of single cell extracellular vesicles secretion. With this platform, we uncovered previously undifferentiated single cell heterogeneity underlying EVs secretion within a phenotypically similar cell population. We then applied the platform to analyze a subgroup of UM-SCC6 cells with high invasion characteristic and observed some EVs expression (e.g. ^CD63+^EVs) decreased in metastatic tumor cells. And the results were further confirmed with OSCC patient samples. These results demonstrated our platform can generate critical information to potentially distinguish and quantitate invasive cell states, which can be used to monitor tumor invasiveness and tailor the therapeutic strategy for individual patient. In addition, these EVs that act as intercellular mediators for cell–cell communication in tumor microenvironment may also be used as new therapeutic targets for personalized medicine.

Tumor microenvironment is collectively shaped by complex signaling networks composed of different mediators, including cytokines, EVs, etc. Direct measurement of different mediators from the same single cells was highly desirable to generate information that inspires deeper understanding of tumor microenvironment and decode the complex signaling network embedded in it. With this platform, we successfully realized multiplexed profiling of two different intercellular communication mediators (5-plxed EVs and 3-plexed proteins) simultaneously from the same single cells, which cannot be obtained using other methods. We observed proteins secretion and EVs secretion were dominated by respective cell subgroups within the population, highlighting the unique advantage associated with multidimensional, multiplexed profiling to resolve the correlation between each parameter. This multidimensional analysis strategy may open up new avenues for uncovering new biology at single cell level.

Notably, the platform is applicable to different cellular types and sources with minimal sample consumption, which makes it especially suitable for rare clinical sample analysis, like circulation tumor cells(44), or fine-needle aspirate (FNA)(18). The proteomic parameters of EVs detection can be further increased if more microchannels are paralleled or spectral encoding is adopted for multi-color detection. When combined with other single cell analysis technologies or different types of perturbations, it could provide more comprehensive information to map the correlation between different functional mediators in cellular microenvironment at different biomimetic models (12, 19, 28, 31). We believe this platform holds great potential to become a broadly applicable tool for in-depth EV analysis in both basic and translational research, like tumor biopsies in precision medicine.

## Methods

### PDMS microchip fabrication

The molds for antibody patterning and single cell capture were fabricated by photolithography with SU8 3035 (Microchem, USA) and treated with TMCS (Trimethylchlorosilane, Sigma-Aldrich) overnight to facilitate peeling PDMS (Polydimethylsiloxane) off the mold. PDMS prepolymer and curing reagent were mixed at 10:1 ratio (RTV615, Momentive), poured onto the mold and cured in the oven at 80°C for 1 hr. The PDMS microchip was bonded with premium grade microarray glass slide (poly-L-lysine coated, Thermo Fisher) after the inlet and outlet holes were punched out. Then it was baked at 80°C for additional 2hrs to complete thermal bonding. The PDMS microwell array for single cell culture was cleaned with ultra-sonication in ethanol and blown dry before use.

### Flow patterning antibodies barcode glass slide

After the PDMS microchip with high density parallel microchannels were assembled with poly-L-lysine coated glass slide, each antibody (**Table S1)** was pushed through individual microchannels until complete dry with 1psi pressured N_2_. The antibodies barcode glass slide was blocked with 1% BSA (Roche, USA) for 1hr to reduce nonspecific adsorption. Then it was washed with DPBS, 50/50 DPBS/DI water and DI water sequentially. The antibody slide was spun dry in slide centrifuge and stored at 4°C before use.

### Cell culture

Human oral squamous carcinoma cell line (SCC25) (American Type Culture Collection) was cultured in MEM medium (Gibco, Thermo Fisher Scientific) with 10% fetal bovine serum (FBS, Gibco, Invitrogen), 1% antibiotics (100 U/ml of penicillin G sodium, 100 U/ml of streptomycin) and 1% MEM Non-Essential Amino Acid (Life Technologies). FBS was ultra-centrifuged at 100,000 x g at 4°C for 4 hours to deplete exosome in it. The Human oral squamous carcinoma cell line (UM-SCC6) (a kind gift from Prof. Songling Wang (Capital Medical University, China)) was cultured in DMEM/HIGH GLUCOSE (HyClone) medium with similar conditions for SCC25. The cells were detached with 0.25% trypsin-0.02% EDTA for 4min, centrifuged at 1000 rpm for 5min, washed and re-suspended in fresh medium before use.

### Isolation of invasion front cells from UM-SCC6

The matrix channel of the isolation microchip was firstly loaded with Matrigel™ (Corning). UM-SCC6 cells were seeded into the cell culture channel in serum-free medium. Then cell culture medium containing 20% FBS was introduced into the stimulation channel. Cells that invaded through the matrix channel and migrated into the stimulation channel were termed as the invasion front cells. These invasion front cells were collected by trypsinization and expanded to repeat the above mentioned steps to generate second round of invasion front cells. The third round of invasion front cells of UM-SCC6 were repeated again and collected as UM-SCC6M cells.

### OSCC patient tissue samples

Human OSCC patient samples were obtained from the Affiliated Hospital of Dalian Medical University. The collection and use of human samples was approved by the Ethics Committee of Dalian Medical University. Patient primary tissue was firstly minced with an ophthalmic surgical scissors to approximately 1mm^3^ pieces and then pipetted repeatedly with DPBS containing 2% antibiotics. The tissue was then detached with 0.25% trypsin-0.02% EDTA for 20-40min at 37°C, shaking once every 5min. The tissue was then detached with collagenase I on the shaker until the tissue became flocculent. The tube containing the flocculent tissue was placed in a 37°C, 5% CO_2_ incubator for 5 min. The following flocculent precipitate was then spread evenly across the culture dish coated with collagenase I. The culture dish was placed in a 37° C, 5% CO_2_ incubator for 1hr and the culture medium DMEM-HG was added dropwise. Change the medium periodically until the cells became confluent in the culture dish. Then the cells were detached with 0.25% trypsin-0.02% EDTA and re-suspended in fresh medium for experiment.

### Single cell EVs secretion analysis procedures

The PDMS microchambers array for single cell assay was treated with O_2_ plasma (Harrick Plasma PDC-32G) for 1min before single cell experiment and blocked with cell culture medium (with 10% FBS) to maintain surface hydrophilic, which will facilitate cell loading and minimize nonspecific protein adsorption. Sample cells were pre-stained with cell viability dye Calein-AM green at 37°C for 30min and re-suspended into fresh medium at defined density. The cells were then pipetted onto microchambers array at 2×10^5^cells/mL cell density, 200μL per chip. After cells settled down into microchambers within 5 minutes, antibodies barcode glass slide was imposed onto the top of microchambers array and clamped together to trap single cells. The microchip trapped with single cells was imaged with a Nikon Eclipse TiE microscope with an automatic stage to record the cell number/position information. The clamp was removed after overnight incubation to finish detection procedures. The glass slide was incubated with a cocktail of detection antibodies (biotin-IL-8, biotin-IL-6, biotin-MCP-1, biotin-CD63) for 1 hour and stained with streptavidin-APC or streptavidin-PE (eBioscience, 1:100 dilution) for another 30min. Then it was washed thoroughly with DPBS, 50/50 DPBS/DI water and DI water sequentially, the glass slide was spun dry and scanned with a GenePix 4300A fluorescence scanner (Molecular Devices).

## Data analysis

The images for single cell counting (bright field and fluorescence) can be processed in Nikon software (NIS-Elements Ar Microscope Imaging Software) by defining threshold in combined images to realize automated cell counting. The fluorescence detection image was analyzed with GenePix Pro software (Molecular Devices) by creating and aligning the microchambers array template followed by extraction of mean fluorescence intensity (MFI). The cell counts and corresponding fluorescent data would be matched and processed in Excel (Microsoft) and Graphpad Prism. The thresholds to determine positive secretion events were defined as mean + 2 ×SD of zero-cell data. Heatmaps and unsupervised clustering were generated with software Cluster/ Treeview (Eisen Laboratory). viSNE (Dana Pe’er lab)was used to transform complex multi-parameter data into two dimensional categorized maps.

## Supporting Information

This article contains supporting information online.

## Acknowledgements

The project was supported by National Natural Science Foundation of China (Grant No. 21874133, 21605143), Youth Innovation Promotion Association CAS (Grant No. 2018217), and funds from Dalian Institute of Chemical Physics (Grant No. SZ201601).

**Author contributions**

Y.L., T.J.L. and Y.H.J. and designed the research; Y.H.J., L.M.L., H.R.S. performed experiments, D.Y.Q., B.S., F.Y.Z. contributed clinical samples; X.J.L. conducted UM-SCC6M cell isolation; Y.H.J., L.M.L., H.R.S., Y.L., B.C.L. conducted data analyses; Y.L. wrote the manuscript, with extensive inputs from all authors.

**Conflict of interest**

The authors declare no conflict of interest.

## Supplementary Figures

**Figure S1.**
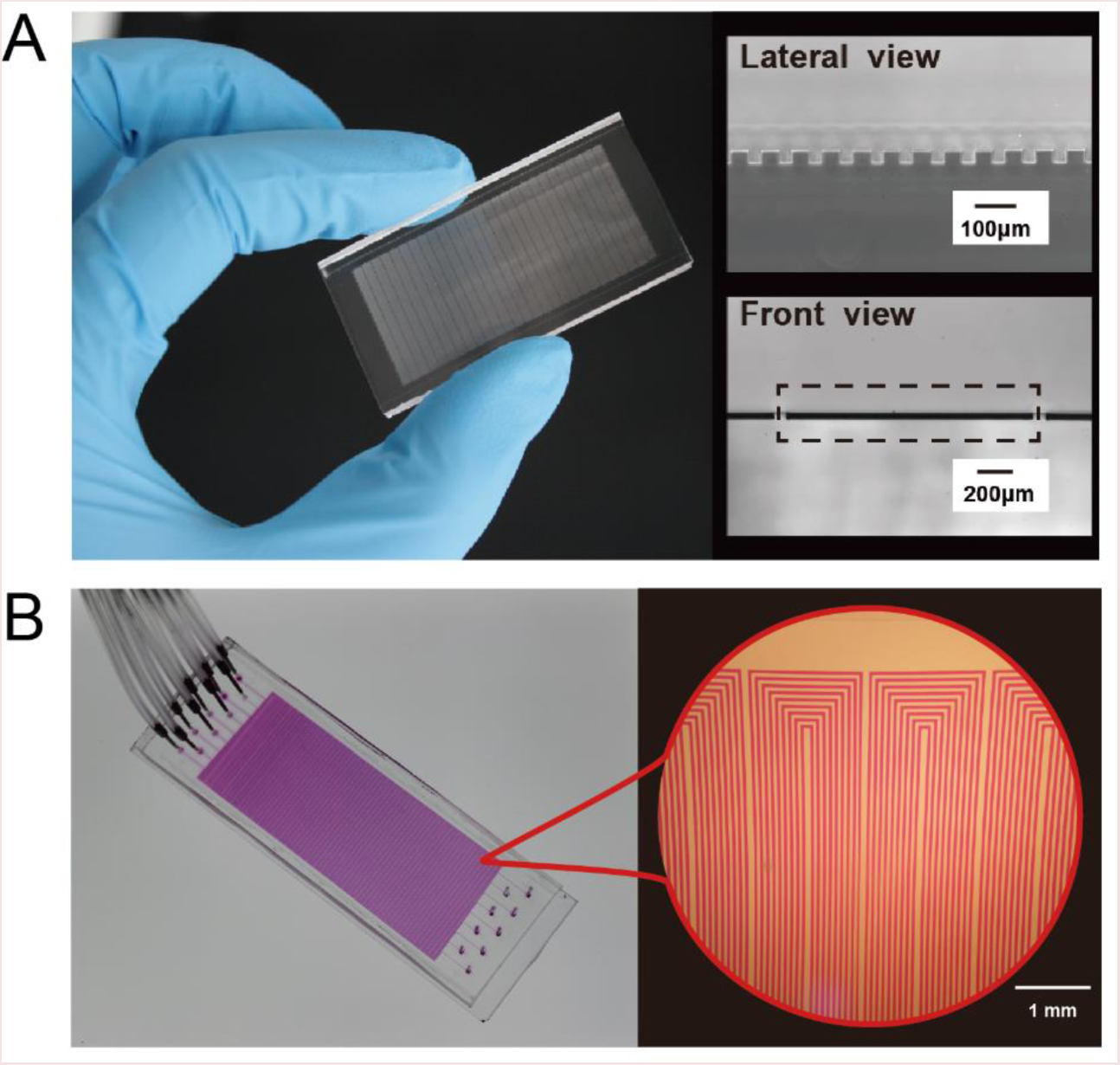
Two functional components of microchip platform for multiplexed profiling of single-cell extracellular vesicles secretion. A, Photographs showing the 6440 PDMS microchambers array to isolate and concentrate EVs secreted from thousands of single cells; B, Images showing the highly parallel microchannel array to pattern spatially resolved antibodies barcode glass slide (the microchannels were filled with red dye solution for visualization).

**Figure S2.**
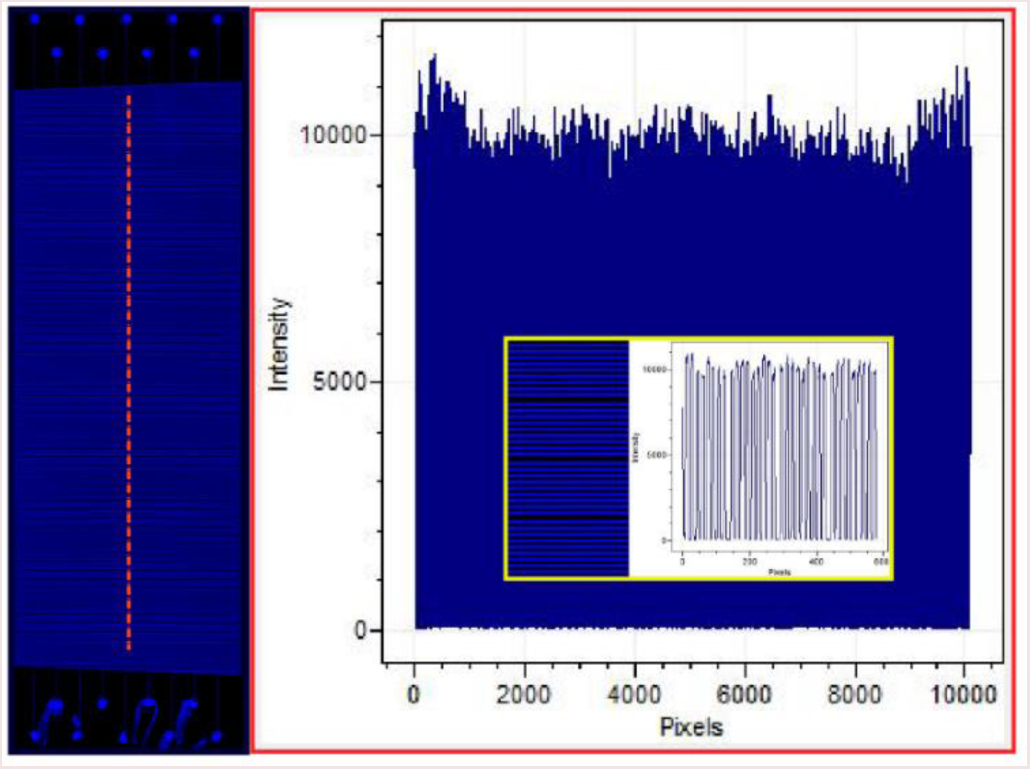
The uniformity characterization of protein patterning on poly-L-lysine glass slide by flow patterning (C.V. <5% across 2 cm x 5.5 cm area). 3μL fluorescently labeled bovine serum albumin (FITC-BSA, 0.25 mg/mL) was pushed through nine parallel microchannels under 1 psi N_2_ until complete dry. After blocking and washing, it was scanned and analyzed by GenePix 4300A and GenePix Pro software (Molecular Devices).

**Figure S3.**
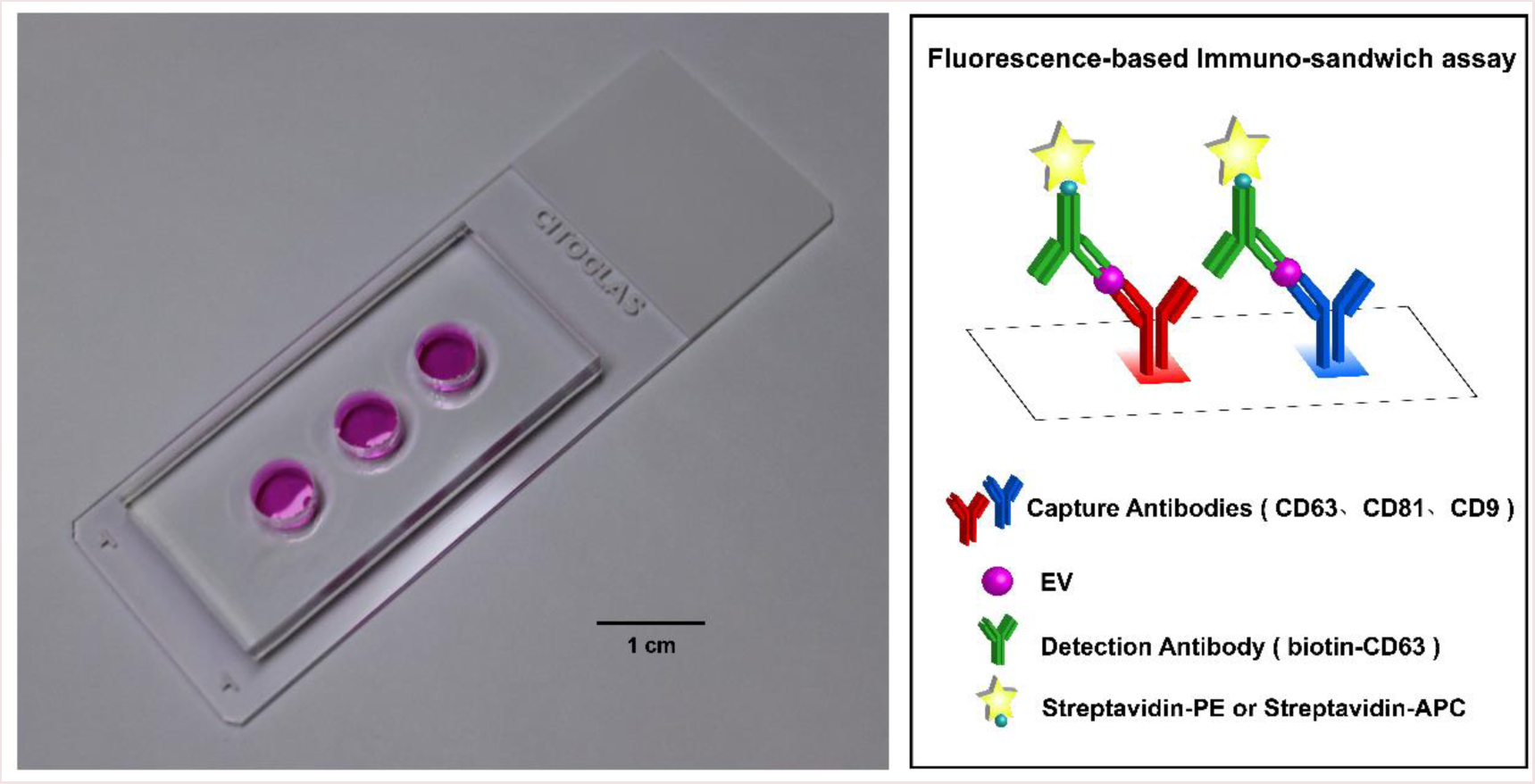
Device and detection principle of cell population EVs secretion assay. Assembly of PDMS microwells slab (diameter=7mm) with poly-L-lysine glass slide for EVs detection with samples from population cells (Left). Schematic illustrating double positive detection strategy based on different epitopes recognition for EVs detection (right).

**Figure S4.**
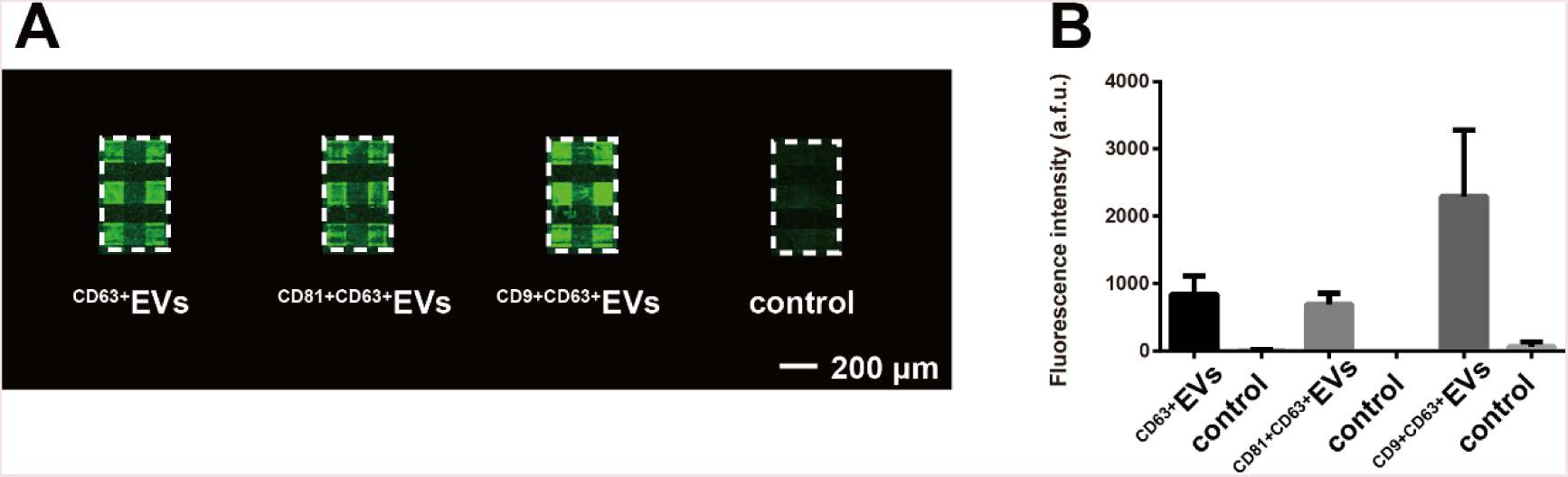
Validation of CD63, CD9, CD81 antibodies barcode to capture/detect EVs from UMSCC6 supernatant. A: Fluorescence images and B: quantification.

**Figure S5.**
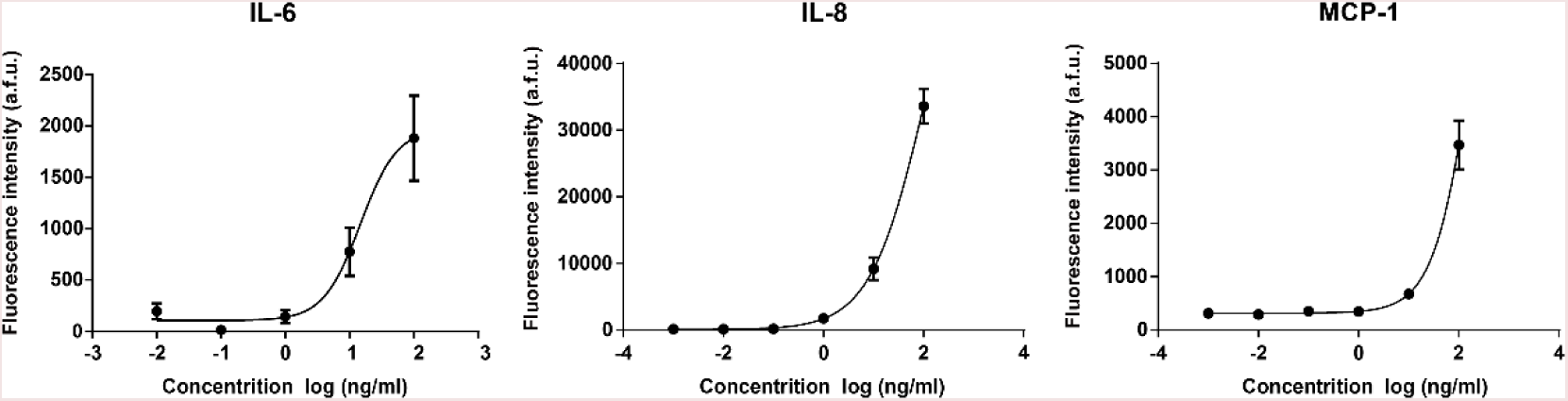
Calibration curves obtained with corresponding recombinant proteins (x, y axes: log-scaled). These titration curves have been fit with a 4PL curve and the fluorescence intensities presented were averaged from 8 spots for each protein.

**Figure S6.**
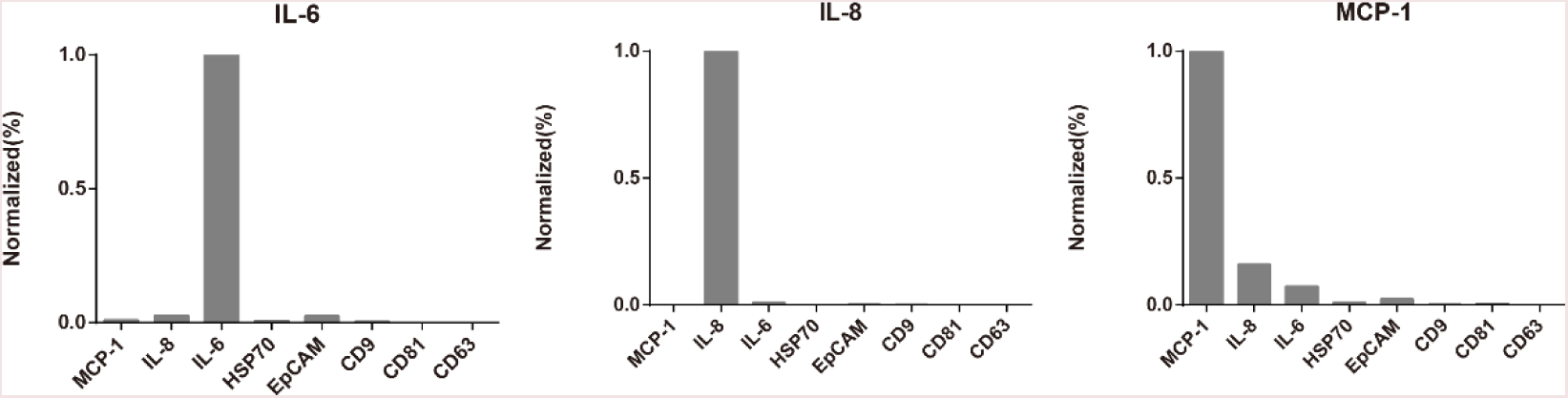
Cross reactivity test. Most of the antibodies used in this study are monoclonal antibodies to ensure good specificity and reduce cross reaction. The test was conducted by spiking a single recombinant protein solution (100 ng/mL) to antibodies barcode containing all capture antibodies, followed by detection with a mixture of detection Abs (IL-6, IL-8, MCP-1 and CD63). The fluorescence intensity was normalized into percentage.

**Figure S7.**
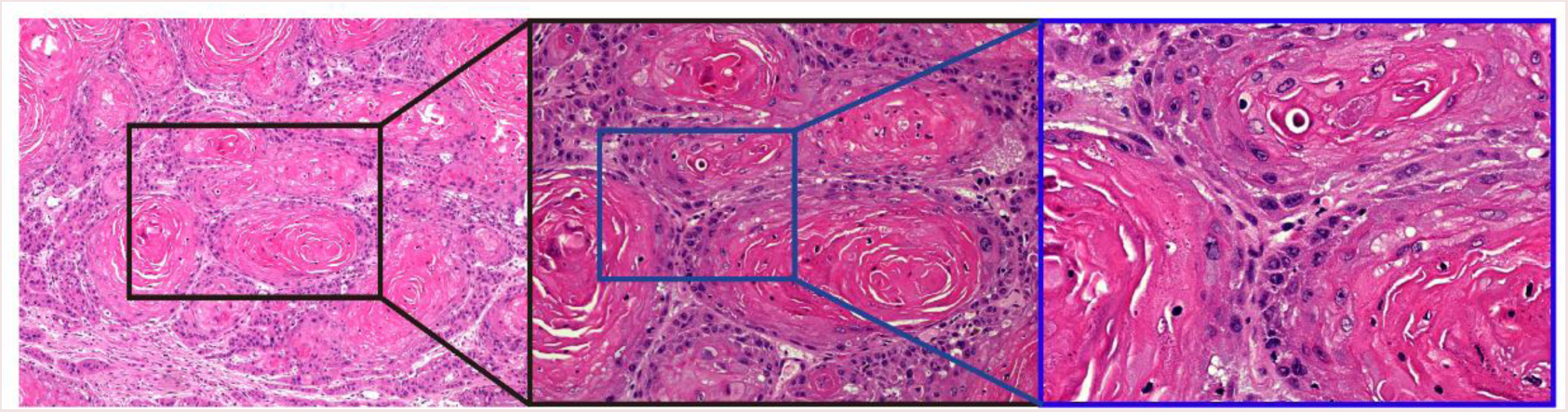
HE staining of patient 3 tissue showed highly differentiated oral squamous cell carcinoma. We observed the intercellular bridges, keratin pearl and a few mitoses, while the nucleus and cell pleomorphic were not obvious.

**Figure S8.**
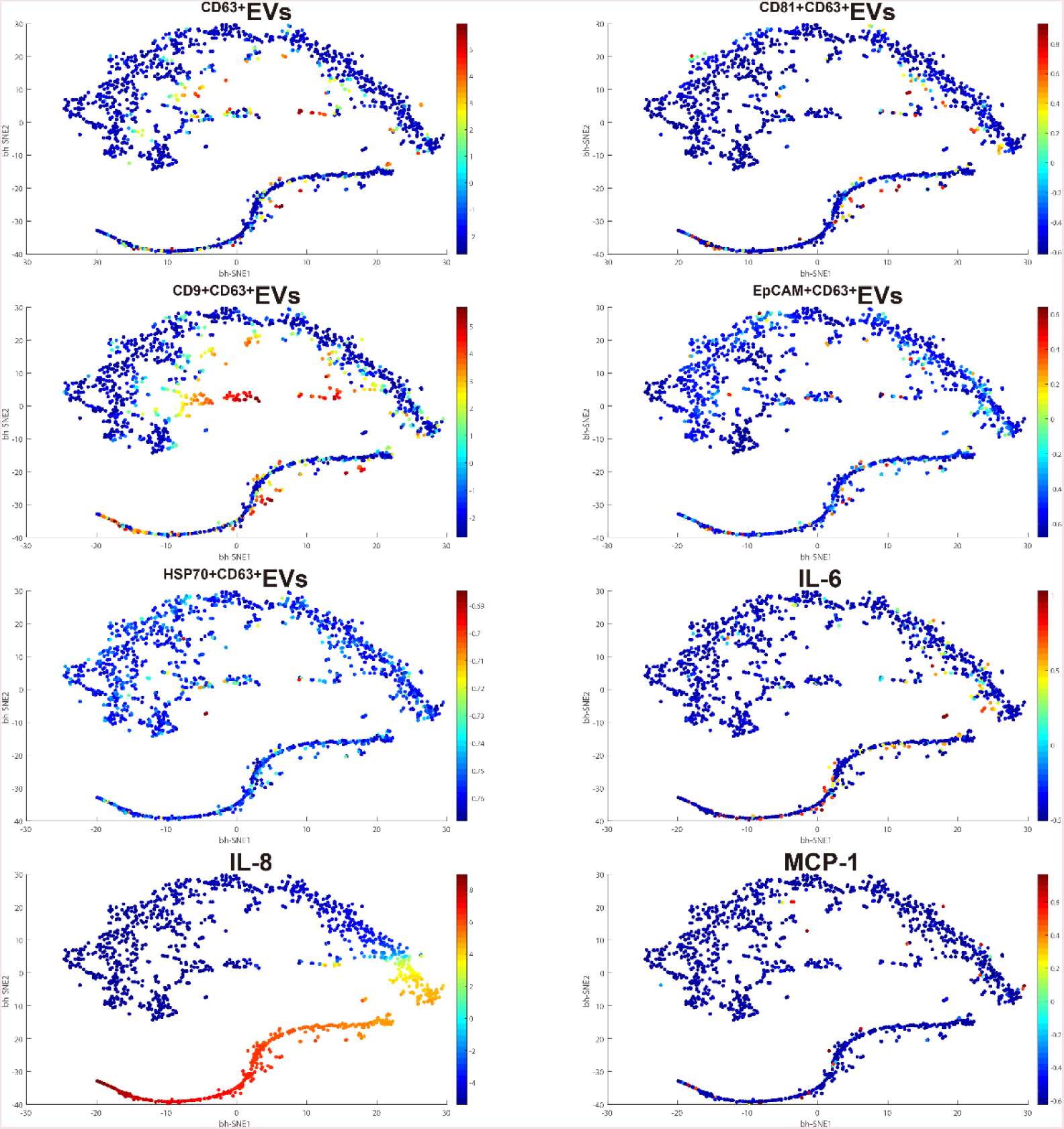
EVs and protein distribution in the viSNE maps for SCC25 single cells.

**Figure S9.**
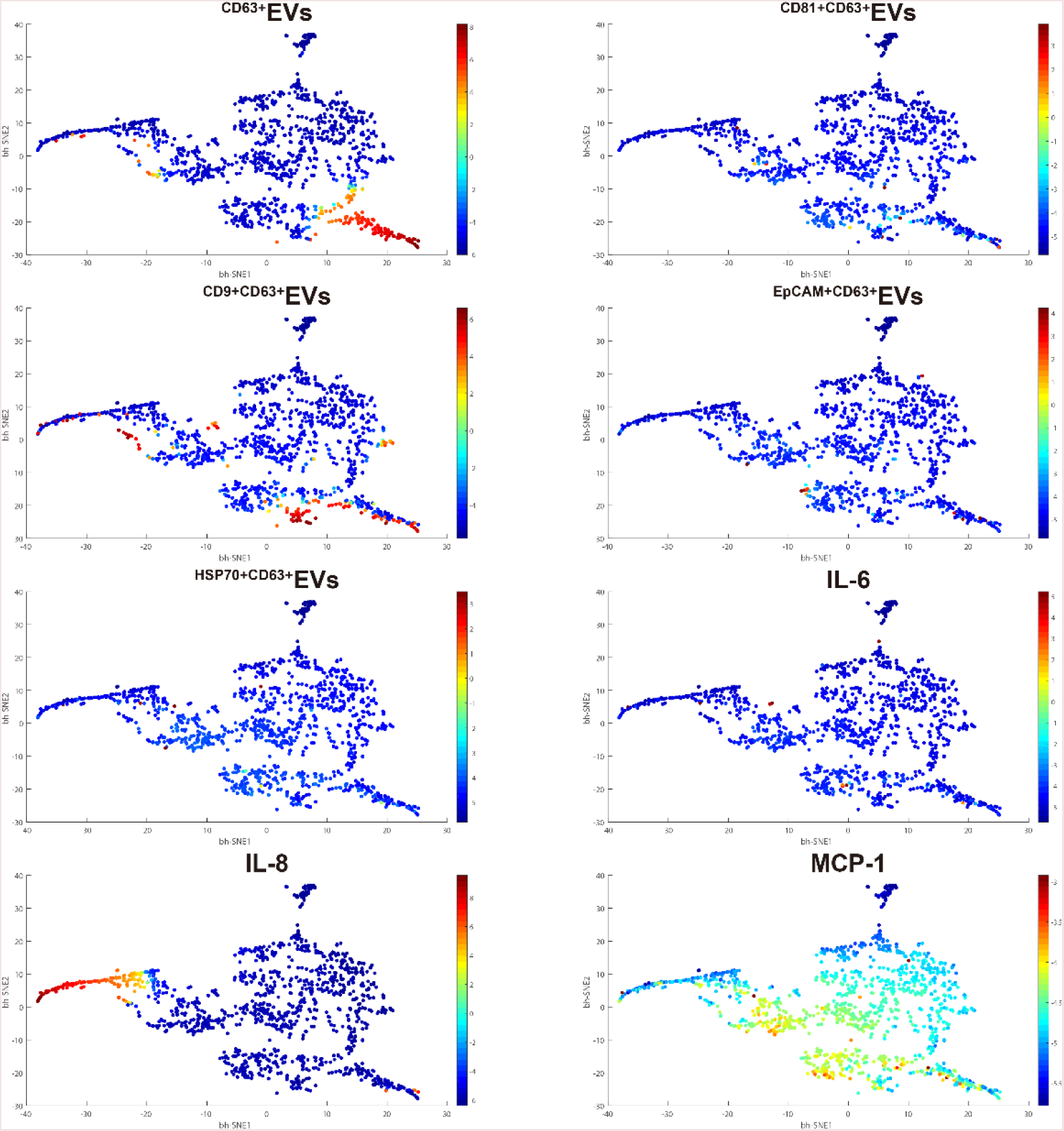
Whole panel secretion distribution in the viSNE maps for UM-SCC6 single cells.

**Figure S10.**
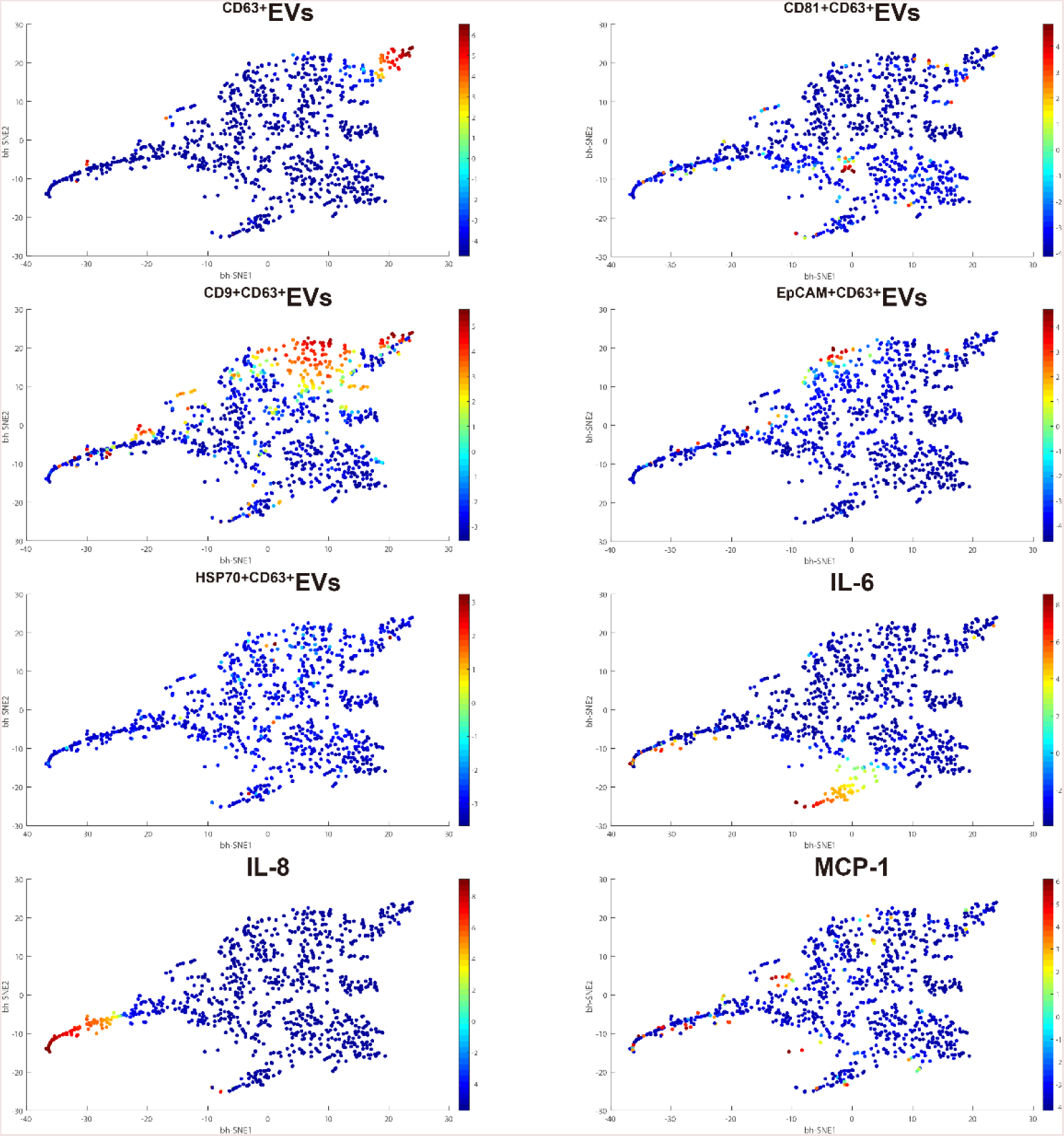
EVs and proteins distribution in the viSNE maps for Patient 1.

**Figure S11.**
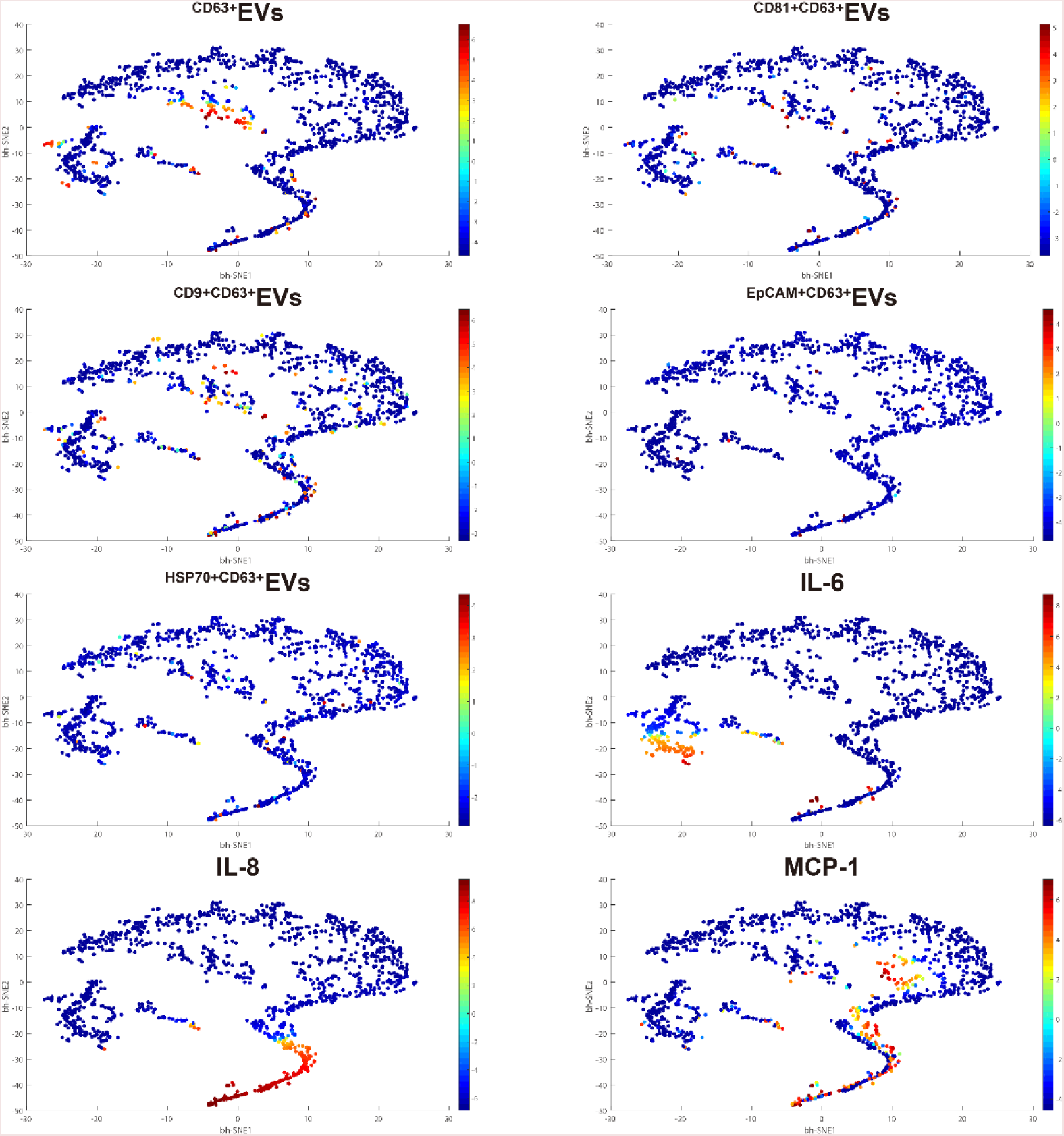
EVs and proteins distribution in the viSNE maps for Patient 2.

**Figure S12.**
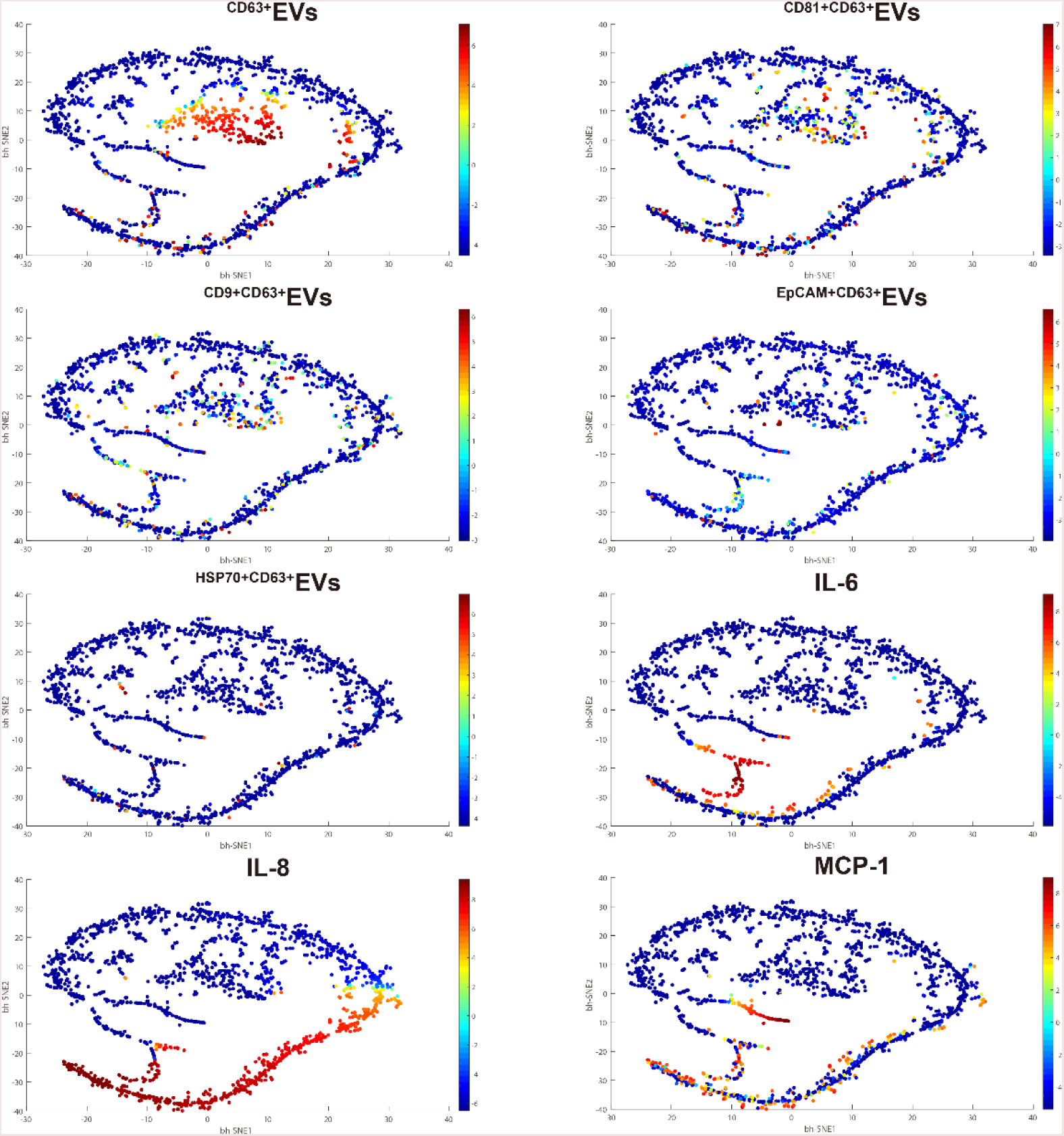
EVs and proteins distribution in the viSNE maps for Patient 3.

**Table S1.**
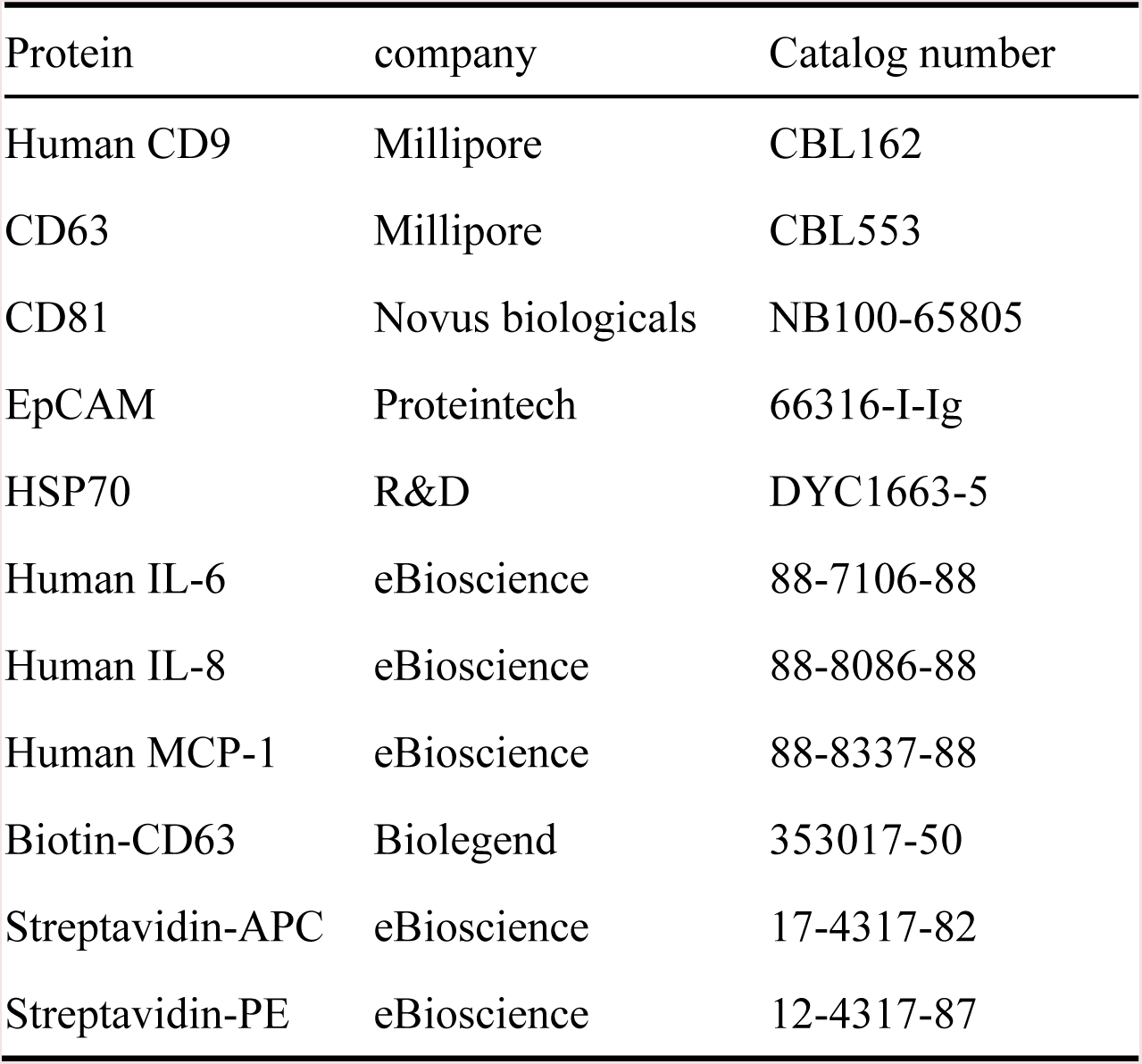
Summary of antibodies used.

**Table S2.** Brief summary of the patients’ medical records.

| Patient | Gender | Age | Tumor type | Location | Differentiation | Metastasis |
| --- | --- | --- | --- | --- | --- | --- |
| 1 | Male | 55 | OSCC | Mouth floor | high | Yes(cervical lymph node) |
| 2 | Male | 38 | OSCC | tongue | high | Yes(submandibular, deep cervical lymph) |
| 3 | Male | 60 | OSCC | tongue | high | No |

## References

1. Raposo G & Stoorvogel W (2013) Extracellular vesicles: Exosomes, microvesicles, and friends. J. Cell Biol. 200(4):373-383.

2. El Andaloussi S, Maeger I, Breakefield XO, & Wood MJA (2013) Extracellular vesicles: biology and emerging therapeutic opportunities. Nature Reviews Drug Discovery 12(5):348-358.

3. van der Pol E, Boing AN, Harrison P, Sturk A, & Nieuwland R (2012) Classification, Functions, and Clinical Relevance of Extracellular Vesicles. Pharmacological Reviews 64(3):676-705.

4. Fevrier B & Raposo G (2004) Exosomes: endosomal-derived vesicles shipping extracellular messages. Curr. Opin. Cell Biol. 16(4):415-421.

5. Gyorgy B, et al. (2011) Membrane vesicles, current state-of-the-art: emerging role of extracellular vesicles. Cell. Mol. Life Sci. 68(16):2667-2688.

6. Lo Cicero A, Stahl PD, & Raposo G (2015) Extracellular vesicles shuffling intercellular messages: for good or for bad. Curr. Opin. Cell Biol. 35:69-77.

7. Muralidharan-Chari V, Clancy JW, Sedgwick A, & D’Souza-Schorey C (2010) Microvesicles: mediators of extracellular communication during cancer progression. J. Cell Sci. 123(10): 1603-1611.

8. Shao H, et al. (2012) Protein typing of circulating microvesicles allows real-time monitoring of glioblastoma therapy. Nature Medicine 18(12): 1835-+.

9. Yang KS, et al. (2017) Multiparametric plasma EV profiling facilitates diagnosis of pancreatic malignancy. Science Translational Medicine 9(391).

10. Tian Y, et al. (2018) Protein Profiling and Sizing of Extracellular Vesicles from Colorectal Cancer Patients via Flow Cytometry. Acs Nano 12(1):671-680.

11. Zabeo D, et al. (2017) Exosomes purified from a single cell type have diverse morphology. Journal of extracellular vesicles 6(1):1329476-1329476.

12. Shao H, et al. (2018) New Technologies for Analysis of Extracellular Vesicles. Chemical Reviews 118(4):1917-1950.

13. Shurtleff MJ, et al. (2017) Broad role for YBX1 in defining the small noncoding RNA composition of exosomes. Proceedings of the National Academy of Sciences of the United States of America 114(43):E8987-E8995.

14. Lotvall J, et al. (2014) Minimal experimental requirements for definition of extracellular vesicles and their functions: a position statement from the International Society for Extracellular Vesicles. Journal of extracellular vesicles 3:26913-26913.

15. Kowal J, et al. (2016) Proteomic comparison defines novel markers to characterize heterogeneous populations of extracellular vesicle subtypes. Proceedings of the National Academy of Sciences of the United States of America 113(8):E968-E977.

16. Yoshioka Y, et al. (2014) Ultra-sensitive liquid biopsy of circulating extracellular vesicles using ExoScreen. Nat. Commun. 5:8.

17. Lee K, et al. (2018) Multiplexed Profiling of Single Extracellular Vesicles. Acs Nano 12(1):494-503.

18. Ullal AV, et al. (2014) Cancer Cell Profiling by Barcoding Allows Multiplexed Protein Analysis in Fine-Needle Aspirates. Science Translational Medicine 6(219):219ra219-219ra219.

19. Heath JR, Ribas A, & Mischel PS (2016) Single-cell analysis tools for drug discovery and development. Nature Reviews Drug Discovery 15(3):204-216.

20. Wang DJ & Bodovitz S (2010) Single cell analysis: the new frontier in ‘omics’. Trends Biotechnol. 28(6):281-290.

21. Colombo M, et al. (2013) Analysis of ESCRT functions in exosome biogenesis, composition and secretion highlights the heterogeneity of extracellular vesicles. J. Cell Sci. 126(24):5553-5565.

22. Willms E, et al. (2016) Cells release subpopulations of exosomes with distinct molecular and biological properties. Sci Rep 6:12.

23. Chiu YJ, Cai W, Shih YRV, Lian I, & Lo YH (2016) A Single-Cell Assay for Time Lapse Studies of Exosome Secretion and Cell Behaviors. Small 12(27):3658-3666.

24. Son KJ, et al. (2016) Microfluidic compartments with sensing microbeads for dynamic monitoring of cytokine and exosome release from single cells. Analyst 141(2):679-688.

25. Verweij FJ, et al. (2018) Quantifying exosome secretion from single cells reveals a modulatory role for GPCR signaling. J. Cell Biol. 217(3):1129-1142.

26. Fan R, et al. (2008) Integrated barcode chips for rapid, multiplexed analysis of proteins in microliter quantities of blood. Nature Biotechnology 26(12):1373-1378.

27. Ma C, et al. (2011) A clinical microchip for evaluation of single immune cells reveals high functional heterogeneity in phenotypically similar T cells. Nature Medicine 17(6):738-U133.

28. Shi QH, et al. (2012) Single-cell proteomic chip for profiling intracellular signaling pathways in single tumor cells. Proceedings of the National Academy of Sciences of the United States of America 109(2):419-424.

29. Lu Y, et al. (2013) High-Throughput Secretomic Analysis of Single Cells to Assess Functional Cellular Heterogeneity. Analytical Chemistry 85(4):2548-2556.

30. Xue Q, et al. (2015) Analysis of single-cell cytokine secretion reveals a role for paracrine signaling in coordinating macrophage responses to TLR4 stimulation. Science Signaling 8(381):12.

31. Lu Y, et al. (2015) Highly multiplexed profiling of single-cell effector functions reveals deep functional heterogeneity in response to pathogenic ligands. Proceedings of the National Academy of Sciences of the United States of America 112(7):E607-E615.

32. Xue Q, et al. (2017) Single-cell multiplexed cytokine profiling of CD19 CAR-T cells reveals a diverse landscape of polyfunctional antigen-specific response. Journal for Immunotherapy of Cancer 5.

33. Ma C, et al. (2013) Multifunctional T-cell Analyses to Study Response and Progression in Adoptive Cell Transfer Immunotherapy. Cancer Discovery 3(4):418-429.

34. Rossi J, et al. (2018) Preinfusion polyfunctional anti-CD19 chimeric antigen receptor T cells associate with clinical outcomes in NHL. Blood.

35. Hasina R, et al. (2008) Angiogenic heterogeneity in head and neck squamous cell carcinoma: biological and therapeutic implications. Laboratory Investigation 88(4):342-353.

36. Barile L, et al. (2014) Extracellular vesicles from human cardiac progenitor cells inhibit cardiomyocyte apoptosis and improve cardiac function after myocardial infarction. Cardiovascular Research 103(4):530-541.

37. Li X, et al. (2018) Downregulation of miR-218-5p promotes invasion of oral squamous cell carcinoma cells via activation of CD44-ROCK signaling. Biomedicine & pharmacotherapy = Biomedecine & pharmacotherapie 106:646-654.

38. Jang HI & Lee H (2003) A decrease in the expression of CD63 tetraspanin protein elevates invasive potential of human melanoma cells. Experimental and Molecular Medicine 35(4):317-323.

39. Lupia A, et al. (2014) CD63 Tetraspanin Is a Negative Driver of Epithelial-to-Mesenchymal Transition in Human Melanoma Cells. Journal of Investigative Dermatology 134(12):2947-2956.

40. Lai X, et al. (2017) Decreased expression of CD63 tetraspanin protein predicts elevated malignant potential in human esophageal cancer. Oncology Letters 13(6):4245-4251.

41. Huan J, et al. (2015) Overexpression of CD9 correlates with tumor stage and lymph node metastasis in esophageal squamous cell carcinoma. International Journal of Clinical and Experimental Pathology 8(3):3054-3061.

42. Otero M, et al. (2012) Human Chondrocyte Cultures as Models of Cartilage-Specific Gene Regulation. Human Cell Culture Protocols, eds Mitry RR & Hughes RD (Humana Press, Totowa, NJ), pp 301-336.

43. Amir E-aD, et al. (2013) viSNE enables visualization of high dimensional single-cell data and reveals phenotypic heterogeneity of leukemia. Nature Biotechnology 31(6):545-+.

44. Sinkala E, et al. (2017) Profiling protein expression in circulating tumour cells using microfluidic western blotting. Nat. Commun. 8:14622.

